# Purine nucleotide limitation undermines antibiotic action in clinical *Escherichia coli*

**DOI:** 10.1101/2023.06.22.546106

**Authors:** Paul Lubrano, Thorben Schramm, Elisabeth Lorenz, Alejandra Alvarado, Seraina Carmen Eigenmann, Amelie Stadelmann, Sevvalli Thavapalan, Nils Waffenschmidt, Timo Glatter, Silke Peter, Knut Drescher, Hannes Link

## Abstract

Metabolic variation across pathogenic bacterial strains can impact their susceptibility to antibiotics^1–4^ and promote evolution of antimicrobial resistance (AMR)^5,6^. However, little is known about which metabolic pathways contribute to AMR, and the underlying mechanisms. Here, we measured antibiotic resistance of 15,120 *Escherichia coli* mutants, each with a single amino acid change in one of 346 essential proteins. Most of the mutant strains that showed resistance to either of the two tested antibiotics carried mutations in metabolic genes. Resistance mutations against a β-lactam antibiotic (carbenicillin) were associated with purine nucleotide biosynthesis and limited the supply of ATP. We show that ATP limitation confers both resistance and tolerance against β-lactam antibiotics by upregulating the purine nucleoside transporter PunC. These results are clinically relevant, because an *E. coli* strain isolated from a clinical specimen had a purine nucleotide limitation, which reduced its susceptibility to antibiotics.

## Introduction

Antimicrobial resistance (AMR) is a major threat to global health, and the associated death toll is alarmingly increasing^7^. Among the major contributors to AMR are pathogenic strains of *E. coli*, which have been associated with more than 800,000 deaths^8^. AMR is either a consequence of mobilized resistance genes (e.g. drug-modifying enzymes), or of mutations that change drug-transport or binding to the drug-target^9,10^. Apart from such canonical resistance mechanisms, mutations in genes that are not directly related to the drug or the drug-target can also confer antibiotic resistance or promote its evolution. However, these non-canonical resistance mutations are difficult to identify due to their indirect effect on antibiotic action, which is often mediated by changes in bacterial physiology and metabolism^1,11,12^. For example, mutations in arginine biosynthesis genes of *Mycobacterium smegmatis* upregulated an aminoglycoside-modifying enzyme^2^, and a hypomorphic variant of a CO_2_ metabolism enzyme promoted fluoroquinolone resistance in *Neisseria gonorrhoeae*^3^. Laboratory evolution studies have provided further evidence for the role of metabolism in antibiotic resistance, suggesting that mutations in metabolic genes have clinical relevance^4^, and that they influence the evolutionary pathway towards resistance^5^.

Despite first approaches to map antibiotic resistance mutations at a genome-scale^13^ and with single-nucleotide resolution^14,15^, it remains difficult to delineate metabolic pathways that are most relevant for AMR. Metabolic pathways that are increasingly associated with antibiotic action are purine nucleotide metabolism^16–19^ and respiration^20,21^. Especially ATP levels have been associated with antibiotic tolerance in *E. coli*^22,23^, as well as with persister formation in *Staphylococcus aureus*^24^ and *E. coli*^25,26^. However, how purine nucleotide metabolism interferes with antibiotic action is currently unclear.

Here, we used a massively parallel assay to measure antibiotic resistance of 15,120 *E. coli* mutants, each with a single amino acid change in one of 346 essential proteins. Most mutations that conferred resistance to the β-lactam antibiotic carbenicillin occurred in genes that are involved in purine nucleotide biosynthesis. By following up on three purine mutants (PurA^L75D^, PurM^F105A^ and HisF^V126P^), we found that these strains have bottlenecks in purine biosynthesis, which confer both resistance and tolerance against carbenicillin. We show that purine-mediated resistance and tolerance is due to upregulation of the purine nucleoside transporter PunC under ATP limiting conditions. These results are clinically relevant, because we found an *E. coli* isolate from a clinical specimen with a purine bottleneck, which markedly reduced the killing activity of carbenicillin/sulbactam combination treatment. Together, our results provide a map of antibiotic resistance mutations, show that PunC links purine metabolism to antibiotic action, and highlight the clinical relevance of nutritional conditions for the efficacy of antibiotic treatments.

### Mapping mutations in essential genes that confer antibiotic resistance

We hypothesized that hypomorphic (or partial loss-of-function) mutations in essential genes confer antibiotic resistance because they decrease cellular growth and metabolic activities. To test this hypothesis, we measured antibiotic resistance of an *E. coli* CRISPR library that consisted of 15,120 strains, each with a single amino acid change in one of 346 essential proteins^27^. These mutations were designed such that the amino acid substitution destabilizes the protein, and therefore the mutations are likely to reduce the activity of the gene product.

To identify resistance mutations, we screened the CRISPR library against two antibiotics: carbenicillin as a representative of peptidoglycan targeting β-lactams, and gentamicin as a representative of aminoglycosides that target protein synthesis. The CRISPR library was cultivated on agar plates with minimal glucose medium and carbenicillin or gentamicin at 2X of the minimal inhibitory concentration (MIC). As a reference we cultivated the library on agar plates without carbenicillin or gentamicin (reference-plate). Each experiment was performed twice at different days to test reproducibility (**Fig. 1a**). A non-edited *E. coli* strain carrying the two plasmids for gene editing was used as a control.

**Fig. 1:**
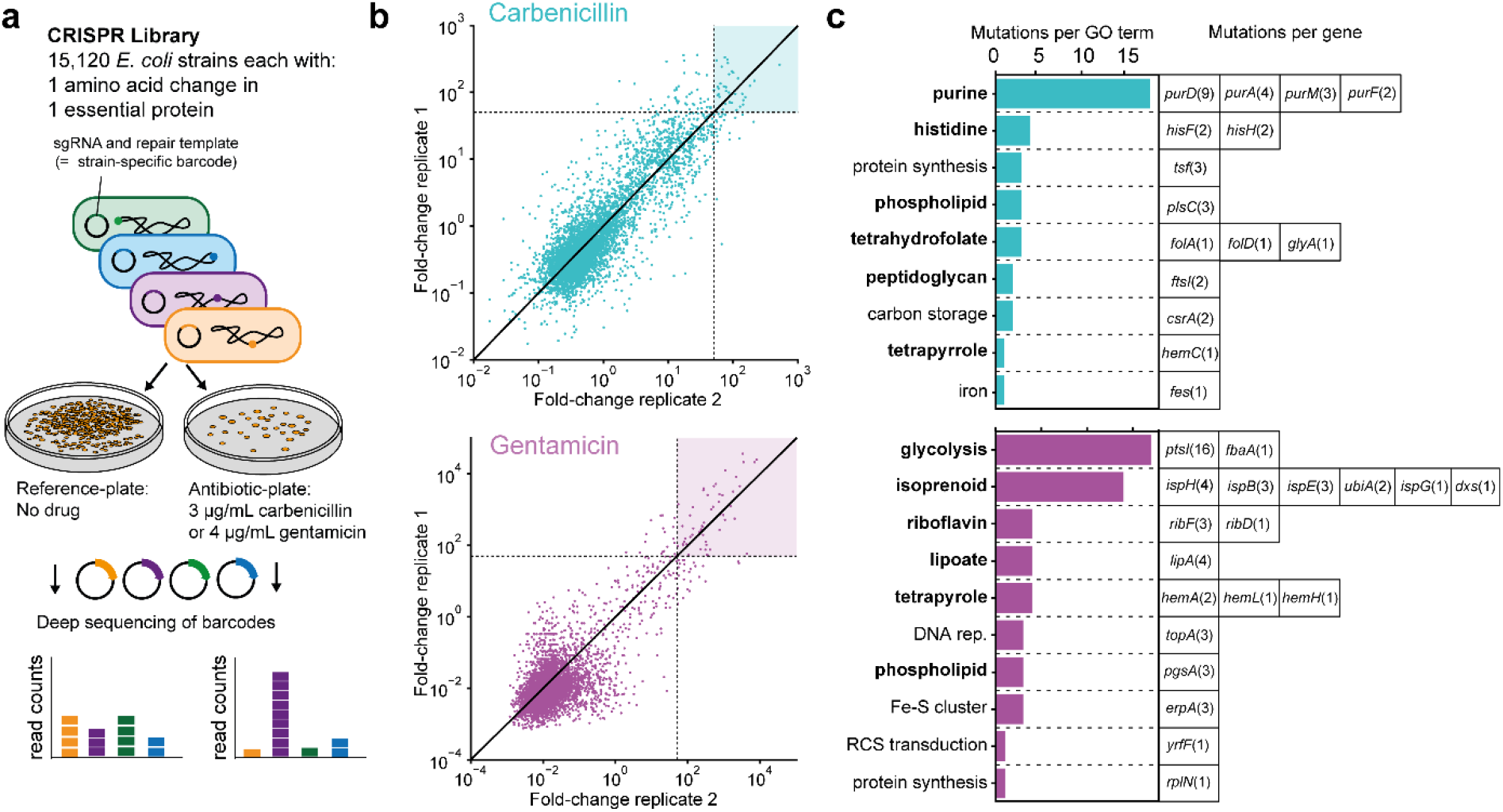
Mapping antibiotic resistance mutations with 15,120 *E. coli* mutants. **a**, Schematic of the CRISPR screen. The CRISPR library included 15,120 *E. coli* strains, each with a single amino acid changes in one of 346 essential proteins. Each mutant has a sgRNA and a repair template (barcode) on a plasmid. The CRISPR library was cultivated on agar plates with and without the antibiotic (n = 2 replicates at different days). Antibiotics were added at 2X of the minimal inhibitory concentration (MIC). Strain-specific barcodes (sgRNA and repair template) were sequenced after 48 h of incubation to determine the composition of the library. **b**, Fold-change of all 15,120 strains in the CRISPR library on carbenicillin (top) and gentamicin (bottom). Fold-changes were calculated by normalizing the barcode abundance on antibiotic-plates by barcode abundance on the reference-plates. Strains with a fold-change >50 in both replicates are considered putatively resistant against the respective antibiotic (blue and magenta region). **c**, Genes with a putative resistance mutation. The number of different resistance mutations per gene is given in brackets. Genes are grouped by gene ontology (GO).

After 48 hours of incubation at 2X MIC, markedly more colonies formed on antibiotic-plates with the CRISPR library compared to antibiotic-plates with the control strain (**Fig. S1**). This was a first indication that the CRISPR library contained resistant mutants. To identify resistant mutants in the CRISPR library, we harvested the colonies from agar plates and determined read counts of strain-specific barcodes (sgRNA and repair templates) by deep sequencing (**Fig. 1a**), which was reproducible between the two replicates (**Fig. S2**). Resistance scores of single mutants were determined as fold-change between sequencing read counts on antibiotic-plates and reference-plates (**Fig. 1b**). Mutants with resistance scores > 50 in both replicates were considered putatively resistant, resulting in 37 resistance mutations for carbenicillin, and 54 resistance mutations for gentamicin (**Fig. 1c**). Out of these 91 mutations, 80 mutations occurred in genes that are involved in metabolism, highlighting the relevance of metabolism for antibiotic action. However, different metabolic processes were associated with gentamicin resistance and carbenicillin resistance, thus indicating that the metabolic states conferring resistance are drug-specific.

In case of the aminoglycoside gentamicin, 16 resistance mutations occurred in *ptsI* which encodes enzyme I of the phosphotransferase system (the major glucose transporter in *E. coli*). Other 14 mutations affected enzymes of the isoprenoid biosynthesis pathway (26%), which is consistent with previous reports about the relevance of isoprenoid metabolism for efficacy of aminoglycoside antibiotics^13^.

In case of the β-lactam antibiotic carbenicillin, 18 out of 37 resistance mutations affected metabolic enzymes in *de novo* biosynthesis of purine nucleotides, and other 4 mutations affected the two subunits of the imidazole glycerol phosphate synthase (HisF and HisG). Although the heterodimer HisFG is an enzyme of the histidine biosynthesis pathway, it also produces the purine intermediate 5-amino-1-(5-phospho-D-ribosyl)imidazole-4-carboxamide (AICAR, **Fig. 2a**). Thus, 22 out of 37 carbenicillin resistance mutations affected purine biosynthesis enzymes, with at least 2 different mutations per gene (**Fig. 1c**). Notably, two resistance mutations occurred in the peptidoglycan DD-transpeptidase encoding gene *ftsI*, which is a direct target of β-lactams ^28^, and the mutations may reduce carbenicillin binding.

**Fig. 2:**
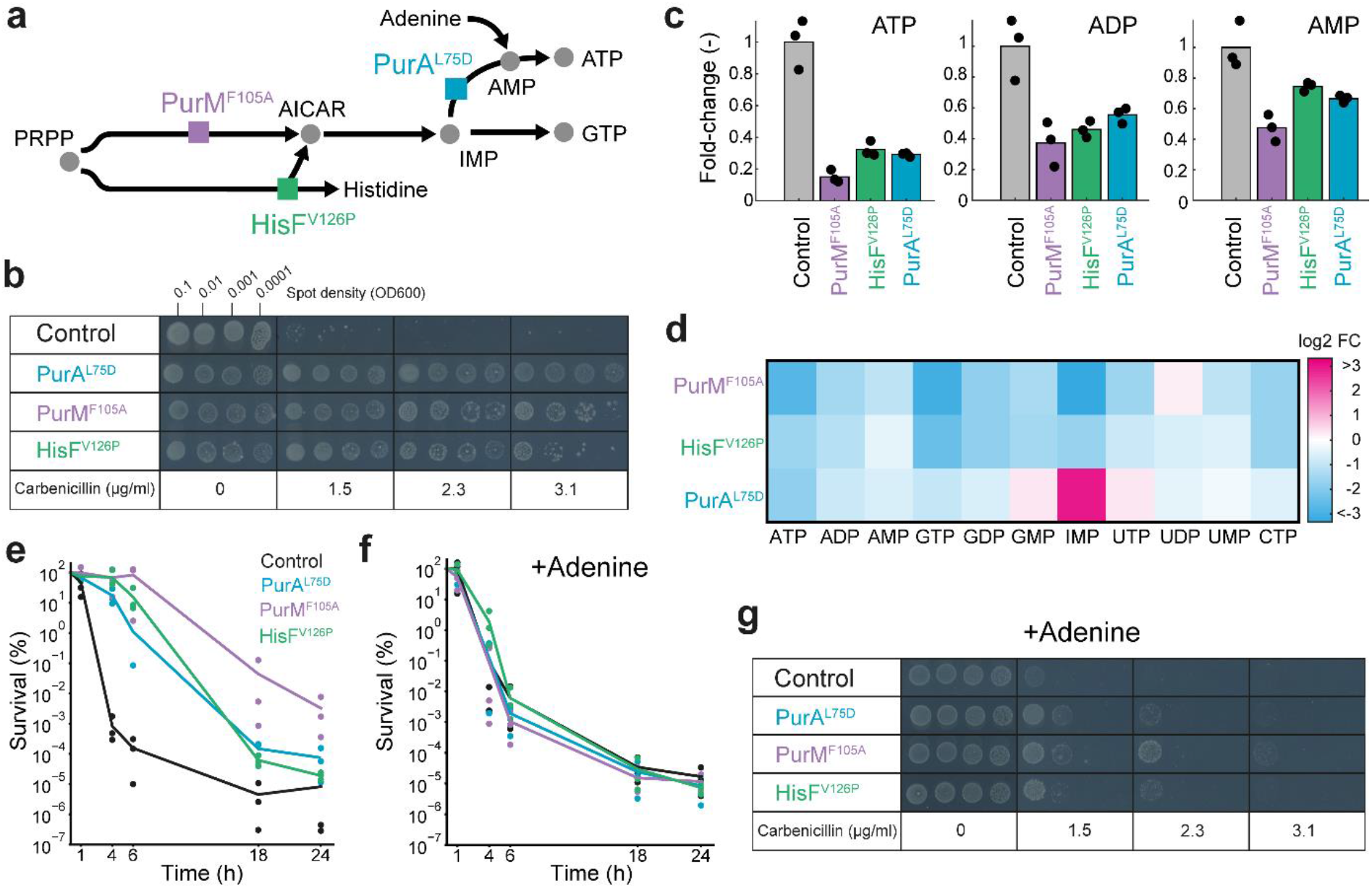
Purine nucleotide limitation confers antibiotic resistance and tolerance. **a**, Schematic of the biosynthesis pathways of purine nucleotides and histidine. Colored boxes indicate the enzymes with resistance mutations: HisF^V126P^, PurM^F105A^ and PurA^L75D^. **b**, Agar dilution assay with the control strain and three re-constructed strains with single mutations (HisF^V126P^, PurM^F105A^ and PurA^L75D^). Each strain was spotted on agar plates with minimal glucose medium containing increasing concentrations of carbenicillin (MIC = 1.5 μg/mL). Multiple inoculum densities were used to assess inoculum effects. Plates were incubated 48 h. Shown is one of n = 2 replicates. **c**, Relative concentration of ATP, ADP and AMP in the HisF^V126P^, PurM^F105A^ and PurA^L75D^ strains. Data are normalized to the control strain and represented as a mean of n = 3 replicates (shown as black dots). **d**, Nucleotide profiles of the HisF^V126P^, PurM^F105A^ and PurA^L75D^ strains. Data are normalized to the control strain and represented as mean of n = 3 replicates. **e**, Time-kill assays with the control strain, and the HisF^V126P^, PurM^F105A^ and PurA^L75D^ strains. Strains were incubated in minimal glucose medium and carbenicillin (50 µg/mL) for the time period indicated on the x-axis. Survival shows colony forming units (CFUs) at the respective time point normalized to CFUs before drug exposure (t= 0 h). **f**, Same time-kill assays as in Fig. 2e with supplementation of adenine. **g**, Same agar dilution assay as in Fig. 2b with supplementation of adenine (shown is one of n = 2 replicates). Note that spot assays were performed on the same plate for one concentration, and scans of plates with different concentrations were assembled into a single figure.

Thus, our CRISPR screen identified putative resistance mutations, which affected predominantly metabolic processes and were drug-specific. This in turn suggests that resistance to antibiotics is not the result of a global growth defect, which has been shown to influence antibiotic lethality^29^, but rather a direct effect of perturbations in specific metabolic pathways.

### Resistance mutations are hypomorphic and introduce metabolic bottlenecks

Next, we validated putative resistance mutations from the CRIPSR screen by re-constructing three carbenicillin resistant mutants: PurM^F105A^ in the upper purine pathway, PurA^L75D^ in the ATP branch, and HisF^V126P^ at the junction of the histidine and the purine pathway (**Fig. 2a**). All three mutants were resistant to carbenicillin, with 2-fold higher MICs than the non-edited control strain (**Fig. 2b**). Additionally, we confirmed gentamicin resistance of the strains PtsI^I330P^, HemA^276Q^, RibD^L364W^, PgsA^V44P^ and IspE^V146W^, which were identified on the gentamicin plates (**Fig. S3**).

To analyze the metabolism of the mutants, we quantified intracellular metabolites during (drug-free) exponential growth on glucose minimal medium by liquid chromatography-tandem mass spectrometry (LC-MS/MS) ^30^. The PtsI^I330P^ strain showed high levels of the glycolysis metabolite phosphoenolpyruvate (PEP, **Fig. S4**), which is the substrate metabolite of the phosphotransferase system (PTS) in *E. coli*. The PTS system is responsible for glucose uptake in *E. coli* and uses PEP to phosphorylate glucose during import. Therefore, we assumed that the gentamicin resistance mutation PtsI^I330P^ decreases the activity of the PTS, which is known to influence antibiotic action in *E. coli* by inducing cyclic-AMP driven gene expression changes in central metabolism^31^.

In the carbenicillin resistant mutants PurA^L75D^, PurM^F105A^ and HisF^V126P^, stronger metabolic changes occurred in nucleotide biosynthesis with decreases between 2- and 5-fold in the purine nucleotide end-products ATP and GTP (**Fig. 2c**). GTP levels were less perturbed in the PurA^L75D^ strain, probably because PurA catalyzes the first step in the ATP branch of the purine pathway. In contrast, PurM^F105A^ and HisF^V126P^ are further upstream in the purine pathway and therefore affected both ATP and GTP levels. The strongest metabolome changes were increases of inosine mono-phosphate (IMP) and 2-(formamido)-N1-(5-phospho-β-D-ribosyl)acetamidine (FGAM) in the PurA^L75D^ and PurM^F105A^ strain, respectively. Because IMP is the substrate metabolite of PurA, and FGAM is the substrate metabolite of PurM, their increase indicates that PurA^L75D^ and PurM^F105A^ are hypomorphic mutations that limit the catalytic capacity of the enzymes. Taken together, decreases of purine end-products (ATP and GTP) and increases of substrate metabolites show that PurA^L75D^, PurM^F105A^ and HisF^V126P^ introduce bottlenecks into the purine pathway. These bottlenecks limit the supply of purine nucleotides, which in turn seems to confer resistance to carbenicillin.

Next, we wondered if purine-mediated carbenicillin resistance is also associated with a higher tolerance against this drug. To test this, we measured time-kill kinetics of the PurA^L75D^, PurM^F105A^ and HisF^V126P^ strains. Indeed, the killing activity of carbenicillin was markedly reduced in the three purine mutants compared to the un-edited control strain (**Fig. 2e**). For example, the PurM^F105A^ strain population remained fully viable during 6 hours of carbenicillin treatment, while less than 0.001% of the control strain population survived this treatment.

Thus, purine mutations confer both resistance and tolerance against carbenicillin. Next, we examined whether the purine limitation is directly responsible for these changes in antibiotic action.

### External supply of adenine restores antibiotic action in purine mutants

To obtain additional evidence that a purine limitation confers carbenicillin resistance and tolerance, we used CRISPR interference (CRISPRi) to decrease the concentration of PurA. Akin to the PurA^L75D^ strain, a strain with CRISPRi-knockdown of *purA* had low ATP levels and high IMP levels (**Fig. S5**), demonstrating that CRISPRi of *purA* introduced a bottleneck in the purine pathway. The *purA*-knockdown was both resistant and tolerant against carbenicillin (**Fig. S5**), which confirmed our hypothesis that a bottleneck in purine nucleotide biosynthesis undermines the efficacy of carbenicillin.

If the purine limitation was responsible for changes in carbenicillin efficacy, we expected that external sources of purine nucleotides would reverse resistance and tolerance of the PurA^L75D^, PurM^F105A^ and HisF^V126P^ strains. *E. coli* can convert external adenine into ATP and GTP by nucleotide salvage pathways that are independent from the *de novo* biosynthesis. As expected, feeding adenine to the PurA^L75D^, PurM^F105A^ and HisF^V126P^ strains reversed their carbenicillin tolerance (**Fig. 2f**) and their carbenicillin resistance (**Fig. 2g**). We also confirmed that adenine supplementation restored the concentration of ATP in the PurA^L75D^ strain to the levels of the control strain (**Fig. S6**). Thus, hypomorphic mutations in genes of the purine pathway cause metabolic bottlenecks that limit the supply of purine nucleotides. This in turn confers resistance and tolerance to carbenicillin.

### The nucleoside transporter PunC is the mechanistic link between purine limitation and antibiotic action

To better understand the molecular consequences of the purine nucleotide limitation in the PurA^L75D^, PurM^F105A^ and HisF^V126P^ strains, we measured their (drug-free) proteomes (**Fig. 3a**). While the PurA^L75D^ and the PurM^F105A^ strains had similar proteome changes, the proteome changes in the HisF^V126P^ strain were unique. The only protein with significant changes in all three strains (log2 fold-change >2 or < -2, p-value <0.05) was the transcription factor PunR (**Fig. 3a**). PunR increased more than 4-fold in the three mutants, which should induce the expression of the nucleoside transporter PunC^32^. Thus, we hypothesized that *E. coli* responds to a purine nucleotide limitation by upregulating PunC, and that increases in the levels of PunC confer resistance and tolerance to carbenicillin (**Fig. 3b**). To test whether PunC is directly responsible for purine-mediated resistance, we created a double mutant with a deletion of *punC* and the PurA^L75D^ mutation (PurA^L75D^Δ*punC* strain) and observed that this mutant was neither resistant nor tolerant to carbenicillin (**Fig. 3c,d**). This implies that PunC is the mechanistic link between the purine limitation in the PurA^L75D^ strain and its carbenicillin resistance and tolerance. Importantly, nucleotide levels in the PurA^L75D^Δ*punC* strain were similar to the PurA^L75D^ strain, and ATP levels remained low (**Fig. 3e**). This rules out the possibility that low ATP levels undermine antibiotic action as a result of the ATP-dependency of antibiotic targets^24^ or ATP driven formation of toxic byproducts^23^. In summary, deletion of *punC* restores antibiotic susceptibility of the PurA^L75D^ strain, thus indicating that upregulation of PunC is the mechanism that confers carbenicillin resistance and tolerance to our purine mutants.

**Fig. 3:**
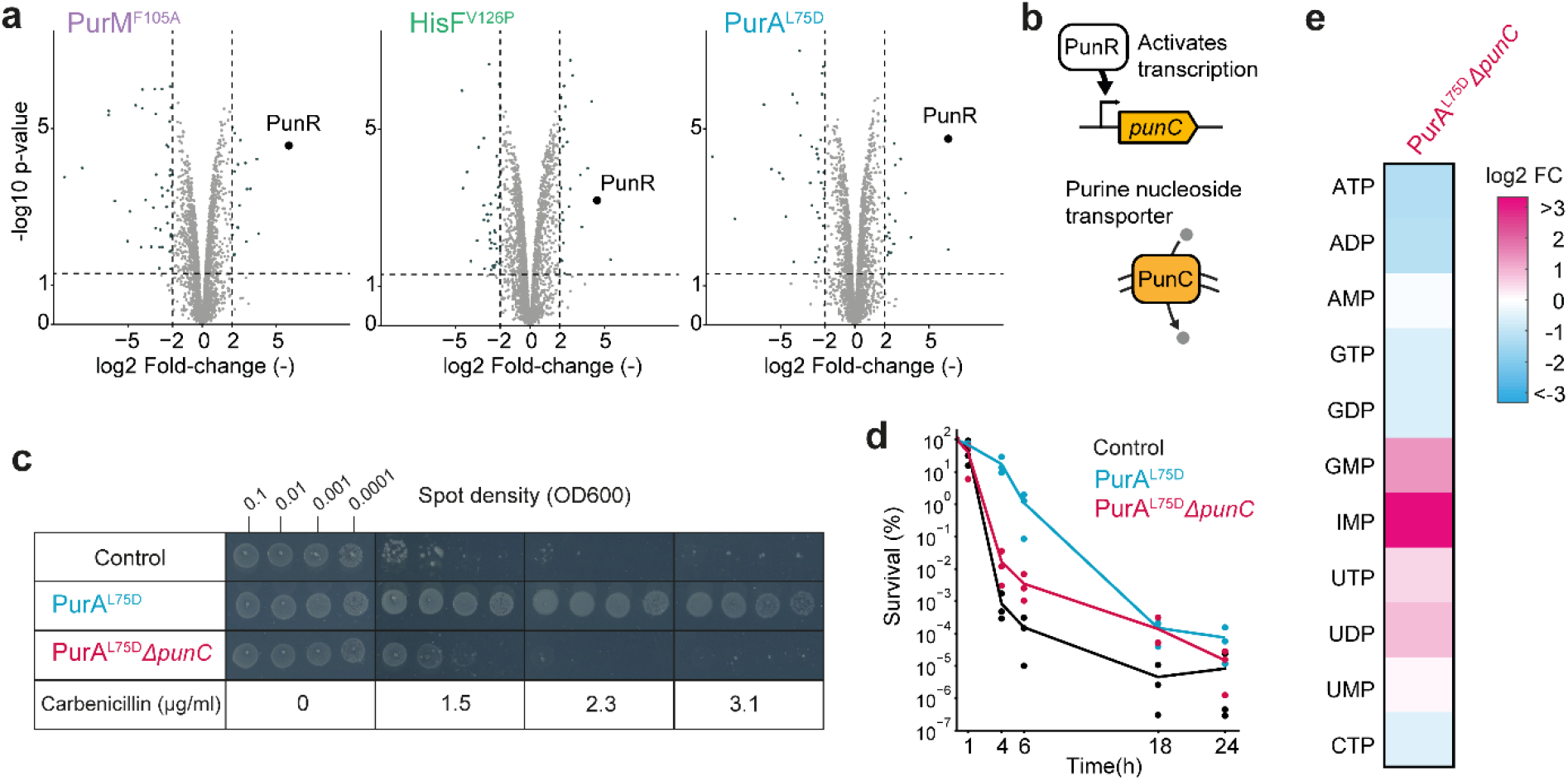
The nucleotide transporter PunC is responsible for purine-mediated antibiotic resistance and tolerance. **a**, Proteome of the HisF^V126P^, PurM^F105A^ and PurA^L75D^ mutants during (drug-free) exponential growth. Shown are fold-changes and p-values of 2511 proteins relative to the control strain. **b**, PunR is a transcription factor that activates the expression of the nucleoside transporter PunC. **c**, Agar dilution assay with the PurA^L75D^ strain and the double mutant PurA^L75D^Δ*punC* (shown is one of n = 2 replicates). **d**, Time-kill assays with the control strain, the PurA^L75D^ strain and the double mutant PurA^L75D^Δ*punC*. Strains were incubated in minimal glucose medium and carbenicillin (50 µg/mL) for the time period indicated on the x-axis. Survival shows colony forming units (CFUs) at the respective time point normalized to CFUs before drug exposure (t= 0 h). **e**, Relative concentrations of intracellular nucleotides in the double mutant PurA^L75D^Δ*punC*. Data are normalized to the control strain and represented as mean of n = 3 replicates. Note that spot assays were performed on the same plate for one concentration, and scans of plates with different concentrations were assembled into a single figure.

### A clinical *E. coli* isolate has a purine nucleotide limitation that confers antibiotic tolerance

Finally, we wondered whether a purine nucleotide limitation is clinically relevant and examined 13 *E. coli* strains that were isolated at the University Hospital of Tübingen from samples of patients with suspected urinary tract infections. All clinical isolates were resistant against at least one class of antibiotics (**Table S1**). To obtain a global picture of the metabolism of the clinical *E. coli* isolates, we measured their metabolomes with flow-injection mass spectrometry^33^ (FI-MS, **Fig. 4a**). The metabolome data indicated a high metabolic variation between the isolates, because the average similarity between the metabolomes was low (**Fig. S7**). Out of 257 measured metabolites, 8 metabolites showed strong increases in at least one isolate (log2 fold-change > 2). The strongest increasing metabolite in this data set was 5-amino-1-(5-phospho-β-D-ribosyl)imidazole (AIR) in isolate #1 (**Fig. 4a**). AIR is an intermediate in the purine biosynthesis pathway and a substrate of 5-(carboxyamino)imidazole ribonucleotide synthase (PurK). We hypothesized that the high AIR levels originated from a metabolic bottleneck at - or near - the PurK catalyzed reaction (**Fig. 4b**). To test whether the supply of purine nucleotides is limited in isolate #1, we cultivated the strain with and without supplementing adenine to the minimal glucose medium. In the absence of adenine, isolate #1 grew poorly compared to another clinical *E. coli* (isolate #6, **Fig. 4c**). Supplementation of 1 mM adenine restored growth of isolate #1, thus supporting our hypothesis that this strain has a bottleneck in purine pathway. To obtain further evidence for the purine limitation in isolate #1, we used targeted LC-MS/MS to measure the concentration of nucleotides and AIR. ATP, ADP and AMP levels in isolate #1 were indeed lower, compared to isolate #6, and the LC-MS/MS data confirmed the high AIR levels detected by FI-MS (**Fig. 4d**). The high levels of AIR, together with low purine end-products suggested that isolate #1 has a metabolic bottleneck in the middle of the purine pathway (between PurK, PurE, PurC, PurB, or PurH, **Fig. 4b**).

**Fig. 4:**
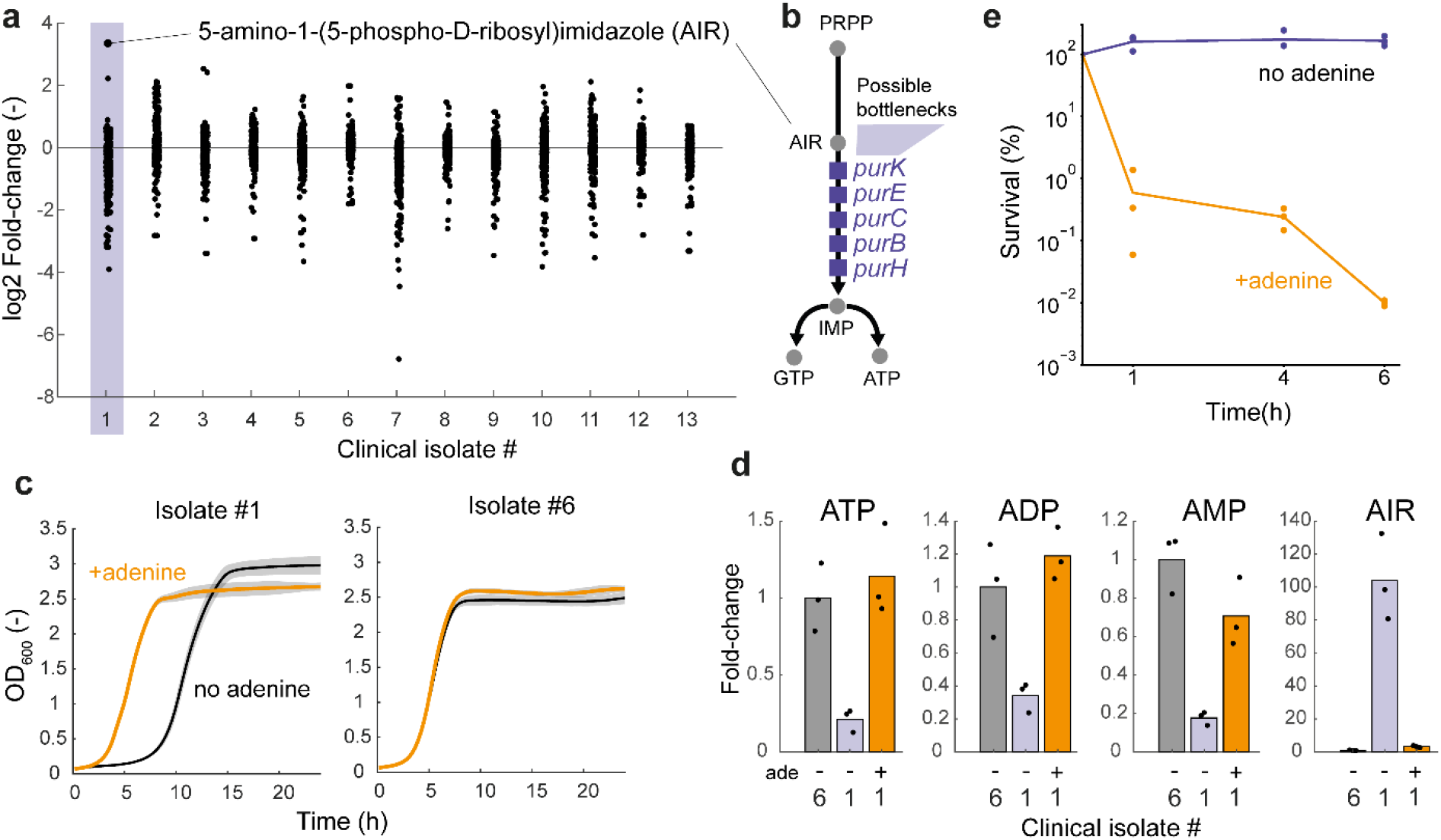
A purine bottleneck undermines carbenicillin/sulbactam treatment of a clinical *E. coli* isolate. **a**, Metabolome of 13 clinical *E. coli* isolates from urine of patients with suspected urinary tract infection. Shown are log2 fold-changes of 257 metabolites relative to the mean of all 13 strains. Dots are means of n = 3 triplicates and only metabolites with a relative standard deviation smaller than 50% are shown. 5-amino-1-(5-phospho-β-D-ribosyl)imidazole (AIR) in isolate #1 is indicated. **b**, Schematic of the purine nucleotide biosynthesis pathway with the location of AIR and the putative bottlenecks. **c**, Growth of isolate #1 and isolate #6 with and without supplementation of adenine. Growth curves show means from n = 3 minimal glucose medium cultures grown in a plate reader. Grey areas show the standard deviation. **d**, Relative concentrations of ATP, ADP, AMP and AIR in isolate #1 (with and without adenine) and isolate #6 (without adenine). Data are normalized to isolate #6 (without adenine) and are represented as mean of n = 3 replicates (shown as black dots). **e**, Time-kill assays with isolate #1 without adenine (violet) and with supplementation of adenine (orange). Isolate #1 was incubated with 100 µg/mL carbenicillin and 12.5 µg/mL sulbactam for the time period indicated on the x-axis. Survival shows colony forming units (CFUs) at the respective time point normalized to CFUs before drug exposure (t = 0 h).

To locate the bottleneck, we sequenced the genomes of isolate #1 and isolate #6 and compared sequences of the suspected purine genes to those of the laboratory *E. coli* strain BW25113. Isolate #1 had 43 mutations in *purK*, eight of which resulted in amino acid changes (**Table S2**). However, seven of the eight amino acid changes occurred both in isolate #1 and isolate #6 and only one amino acid change (PurK^E46G^) was unique to isolate #1. None of the amino acid changes in *purE, purC, purB, purH* was unique to isolate #1 (**Table S2**), thus indicating that PurK^E46G^ is an hypomorphic allele and likely leads to a purine limitation in isolate #1.

Based on the results with our purine mutants, we expected that the purine limitation impairs the efficacy of β-lactam antibiotics in isolate #1. However, isolate #1 was highly resistant against carbenicillin due to a β-lactamase that degrades carbenicillin (**Table S1**). Therefore, we examined the killing activity of a combination treatment of carbenicillin and the β-lactamase inhibitor sulbactam in isolate #1. Consistent with our expectation, carbenicillin/sulbactam exhibited no killing activity in isolate #1 during a 6-hour treatment. However, supplementation of adenine restored killing activity of carbenicillin/sulbactam, such that only 0.01% of the adenine-fed cells survived the 6-hour treatment. Notably, adenine feeding restored ATP levels and AIR levels in isolate #1 (**Fig. 4d**). These data demonstrate that a purine limitation in a clinical *E. coli* isolate impairs the efficacy of carbenicillin/sulbactam, and highlight the clinical relevance of adenine feeding (e.g., by adjuvant therapies that increase adenine levels at the infection site).

## Discussion

In summary, we showed that single hypomorphic mutations can result in metabolic bottlenecks that modulate the concentration of intracellular metabolites and proteins. While such metabolic bottlenecks reduce fitness in the absence of drugs, we found that bottlenecks in glycolysis (PtsI^I330P^) and in purine biosynthesis (PurA^L75D^, PurM^F105A^ and HisF^V126P^) increase fitness in the presence of antibiotics. Purine bottlenecks conferred antibiotic resistance and tolerance by inducing the expression of the purine nucleoside transporter PunC. Previous studies have frequently associated antibiotic efficacy with purine metabolism, either via the concentration of ATP^17,22,24–26^, or adenine and adenosine^18,23^. Hypotheses about why changes in purine metabolism alter antibiotic efficacy include, for example, ATP-dependency of antibiotic targets^24^, or ATP-dependent formation of toxic metabolic byproducts^23^. In the case of our purine mutants we can exclude these hypotheses, because the double mutant PurA^L75D^Δ*punC* had low ATP levels but was susceptible to carbenicillin. Therefore, at least in our purine mutants, the nucleoside transporter PunC mechanistically links purine nucleotide metabolism with antibiotic resistance and tolerance. Future work should clarify whether upregulation of PunC is also the source of the ATP-dependency of antibiotic action observed by others, especially in *S. aureus*^6,24^. Whether PunC is a drug efflux pump or alters antibiotic action by other mechanisms is currently unclear. However, recent data show that PunC is involved in uptake of sulfathiazole and sulfamethoxazole^32^.

Furthermore, it seems possible that changes of transport activities are linked to other metabolism-mediated resistance mechanisms as well. This hypothesis is supported by a recent study in yeast^34^, in which the metabolic adaptations that were necessary to uptake external metabolites decreased intracellular concentrations of antifungal drugs. Thus, future studies could map regulatory metabolites that modulate activity or abundance of transporters, porins and drug efflux pumps, using for instance chemical proteomics^35^. Knowledge about metabolites that actively control antibiotic transport would open up new possibilities for adjuvant therapies to target the nutritional environment at an infection site.

## Acknowledgement

We thank Urs Jenal, Heike Brötz-Oesterhelt and Ilka Bischofs-Pfeifer for discussions. This work was funded by the DFG Cluster of Excellence EXC2124 ‘Controlling Microbes to Fight Infection’ (CMFI). Amplicon sequencing was supported by the Quantitative Biology Center (QBiC), Institute for Medical Genetics and Applied Genomics (IMGAG) and Institute for Medical Microbiology and Hygiene (MGM) of the University of Tübingen.

## Authors contribution

Conceptualization: PL, HL; Methodology: PL, TS, HL; Investigation: all authors; Visualization: PL, HL; Funding acquisition: PL, HL; Project administration: HL; Supervision: HL; Writing – original draft: PL, HL.

## Methods

### Strains

*E. coli* BW25113 was used to construct the CRISPR library and the strains PurA^L75D^, PurM^F105A^, HisF^V126P^. Strain JW1652-1 Δ*punC* from the Keio collection^36^ was used to construct the double mutant Δ*punC* /PurA^L75D^. *E. coli* YYdCas9^37^ was used for CRISPRi. One Shot™ TOP10 *E. coli* (Thermo Fischer #C404010) was used for intermediate cloning.

### Media

Cultivations were performed in LB medium (Sigma #L3522) or M9 minimal medium with glucose as sole carbon source (5 g/L). M9 medium was composed by (per liter): 7.52 g Na_2_HPO_4_ 2 H_2_O, 5 g KH_2_PO_4_, 1.5 g (NH_4_)2_S_O_4_, 0.5 g NaCl. The following components were sterilized separately and then added (per liter of final medium): 1 mL 0.1 M CaCl_2_, 1 mL 1 M MgSO_4_, 0.6 mL 0.1 M FeCl_3_, 2 mL 1.4 mM thiamine-HCl and 10 mL trace salts solution. The trace salts solution contained (per liter): 180 mg ZnSO_4_ 7 H2O, 120 mg CuCl_2_ 2 H_2_O, 120 mg MnSO_4_ H2O, 180 mg CoCl_2_ 6 H_2_O. When needed, M9 agar plates were done by mixing (1:1) a 2X M9 solution with 3 g/L molten agar (Roth #5210.2). In either liquid or agar medium, kanamycin (50 µg/ml; Roth #T832.3) was added when strains harboured a pgRNA plasmid with a kanamycin resistance cassette, and was supplemented with chloramphenicol (30 µg/ml; Merck #C0378-25G) when strains harboured pTS040 and pTS041. When needed, gentamicin (Roth #O233.2) or carbenicillin (Roth #6344.3) were added to the M9 medium at various concentrations specified for each experiment. Adenine (Sigma #A8751-1G) was added to agar and liquid medium at a final concentration of 1 mM, and to agar and liquid medium at final concentration 100 µM for experiments with clinical isolates. To induce dCas9 expression, the CRISPRi experiments were performed with 0.2 µM of anhydrotetracycline (aTc; Cayman Chemicals #10009542) in liquid medium and 1 µM aTc in agar medium.

### Agar dilution assays to measure MICs

M9 agar plates with various concentrations of antibiotics and additives were prepared as described above. Precultures in 4 mL LB were inoculated from glycerol stocks for 8 h at 37°C and 220 RPM shaking, and transferred to M9 medium for overnight incubation at 37°C and 220 RPM. Before starting the assay, precultures were reinoculated and grown in fresh M9 medium to obtain exponentially growing cells. Then, cultures were diluted with fresh M9 medium to set OD600 = 0.1. A 96 well-plate was prepared with three wells containing 135 µL of fresh M9 medium for each strain to be spotted. Each preculture was then 1:10 diluted by mixing 15 µL of the 0.1 OD600 preculture with 135 µL of fresh medium. This process was repeated two times to generate the 1:100 and 1:1000 dilutions. 7 µL of each dilution was then added to the agar plate with a multi-channel pipette to generate the spots. Spots were then left to dry under a flame and plates were incubated for 48h at 37°C. After incubation, plates were imaged with an Epson V370 scanner.

### Cultivation conditions for metabolome and proteome sampling

Precultures in 4 mL LB were inoculated from glycerol stocks for 8h at 37°C 220 rpm shaking and transferred to M9 medium for overnight incubation at 37°C and 220 RPM. M9 pre-cultures in exponential phase were used to inoculate shake flasks containing 10 mL of M9 medium with a starting OD600 of 0.1. Strains were cultivated in triplicates until OD600 reached 0.25-0.6. For metabolomics, flasks were then rapidly transferred to a thermostatically controlled hood at 37°C and an equivalent of OD600 = 1 was sampled. For proteomics, an equivalent of OD600 = 0.8 was transferred to pre-chilled 2 mL tubes.

### Metabolomics measurements

Cultivations were performed as described above. Culture aliquots were vacuum-filtered on a 0.45 μm pore size filter (Merck Millipore #HVLP02500). Filters were immediately transferred into a 40:40:20 (v-%) acetonitrile (Honeywell # 14261-1l)/methanol (VWR # 83638.320)/water extraction solution at -20°C. Filters were incubated in the extraction solution for at least 30 minutes. Subsequently, metabolite extracts were centrifuged for 15 minutes at 13,000 rpm at -9°C and the supernatants were stored at -80°C until analysis.

For targeted metabolomics, metabolite extracts were mixed with a ^13^C-labeled internal standard in a 1:1 ratio. LC-MS/MS analysis was performed with an Agilent 6495 triple quadrupole mass spectrometer (Agilent Technologies) as described previously^30^. An Agilent 1290 Infinity II UHPLC system (Agilent Technologies) was used for liquid chromatography. Temperature of the column oven was 30°C, and the injection volume was 3 μL. LC solvents in channel A were either water with 10 mM ammonium formate and 0.1% formic acid (v/v) (for acidic conditions), or water with 10 mM ammonium carbonate and 0.2% ammonium hydroxide (for basic conditions). LC solvents in channel B were either acetonitrile with 0.1% formic acid (v/v) (for acidic conditions) or acetonitrile without additive (for basic conditions). LC columns were an Acquity BEH Amide (30 × 2.1 mm, 1.7 μm) for acidic conditions, and an iHILIC-Fusion(P) (50 × 2.1 mm, 5 μm) for basic conditions. The gradient for basic and acidic conditions was: 0 min 90% B; 1.3 min 40 % B; 1.5 min 40 % B; 1.7 min 90 % B; 2 min 90 % B. The ratio of ^12^C and ^13^C peak heights was used to quantify metabolites.

For flow-injection metabolomics, metabolite extracts were directly injected into an Agilent 6546 Series quadrupole time-of-flight mass spectrometer (Agilent Technologies, USA) as described previously^33^. The electrospray source was operated in negative and positive ionization mode. The mobile phase was 60:40 isopropanol:water buffered with 10 mM ammonium carbonate (NH4)2CO3 and 0.04 % (v/v) ammonium hydroxide for both ionization modes, and the flow rate was 0.15 mL/min. For online mass axis correction, 2-propanol (in the mobile phase) and HP-921 were used for negative mode and purine and HP-921 were used for positive mode. Mass spectra were recorded in profile mode from 50 to 1700 m/z with a frequency of 1.4 spectra/s for 0.5 min using 10 Ghz resolving power.

### Screening of antibiotic resistance of the CRISPR library

Antibiotic resistance was screened in a pooled CRISPR library with 15,120 *E. coli* mutants that was constructed previously^27^. The CRISPR library and the control strain were each cultivated in 10 mL LB medium at 30°C until OD600 reached 0.5. An equivalent of OD600 = 5 was then centrifuged at 30°C and pellets were resuspended in 10 mL of fresh M9 medium. Centrifugation was repeated to remove traces of LB medium. Cells were resuspended in 10 mL of M9 medium and further incubated at 30°C for 1h. OD600 was then set to 0.5 and 500 µL were used to inoculate 150×20 mm M9 agar plates. Plates were then incubated at 37°C for 48h. After incubation, plates were imaged with an Epson V370 scanner. Then, colonies were harvested from each plate using 7.5 mL of LB medium. OD600 was measured and an equivalent of OD600 = 10 was pelleted in a microcentrifuge tube. Plasmids were then purified by miniprep (Thermo scientist, GeneJET Plasmid Miniprep Kit) for amplification of repair templates and sgRNA (barcodes). Hereafter, 3 ng of plasmid DNA was used for amplification (15 cycles) of the barcodes using two primers suited for further indexing PCRs:

forward primer

5’-TCGTCGGCAGCGTCAGATGTGTATAAGAGACAGGTATCACGAGGCAGATCCTCTG’-3’

reverse primer:

5’-GTCTCGTGGGCTCGGAGATGTGTATAAGAGACAGACTCGGTGCCACTTTTTCAAGTT-3’

Amplicons were purified by AMPure XP PCR beads (Beckman Coulter, #A63881). Using standard Illumina indexing primers, amplicons were indexed in a second PCR and again purified by bead-clean up. Amplicons were pooled and sequenced on an Illumina NextSeq 500 (paired-end, NextSeq™ 500 Mid Output Kit v2.5, #20024908, 300 cycles). Two cartridges were required to yield the desired sequencing depth of around 4 million reads per sample.

### Illumina sequencing data analysis

Demultiplexed paired-end reads were aligned, merged (based on overlapping sequences), and trimmed to the region of interest using a custom Matlab script. The resulting processed reads were mapped against the designed sequences of the library. For each library member, the number of matching reads was counted. Only reads that shared a 100% identity with a designed sequence were considered for further analysis since mutations could indicate a malfunction of the CRISPR-Cas9 genome editing system with no genomic edit. Read counts of 0 were set to 1, to avoid division by 0. Read counts lower than 15 on both reference plates and antibiotic plates were not considered in the analysis. Read counts were normalized by dividing the read counts of each mutants by the total number of reads in a given sample. Fold-changes were calculated by dividing normalized read counts of each mutants on the antibiotic plates by normalized read counts on the reference plates obtained from the same experiment.

### Construction of single CRISPR strains

The PurA^L75D^, PurM^F105A^ and HisF^V126P^ strains were re-constructed with the same method as the CRISPR library. Plasmid pT0S41 was first transformed with electroporation into WT BW25113. Plasmids pTS040 were built by assembling the pTS040 backbone with oligonucleotides (Twist Bioscience) encoding sgRNA and homology arms associated with the desired mutations. Then, plasmids pTS040 were transformed with electroporation after 30 min induction with 7.5 g/L arabinose (lambda red expression). Strains were cultivated for 1 h in SOC medium with kanamycin and 1 µM aTc to induce Cas9 expression. Strains were then plated on LB agar with kanamycin, chloramphenicol and 1 µM aTc. Incubation was done at 37°C overnight. Subsequently, single colonies were picked for colony PCR to amplify the potentially mutated genes of interest. PCR amplicons were purified (Macherey-Nagel #740609) and sent for Sanger sequencing (Eurofins genomics). Sequences were analysed with the Benchling software and the MAFFT algorithm^38^. Strains with the correct mutations were cultivated overnight in 4 mL LB with kanamycin and chloramphenicol to prepare glycerol stocks.

Mutants PtsI^I330P^, RibD^L364W^, PgsA^V44P^, IspE^V146W^ and HemA^L276Q^ were isolated directly from the CRISPR library after gentamicin challenge during a pre-experiment. The library was screened as described above and plated on M9 agar with 4 µg/mL gentamicin. Incubation was done over 48 h at 37°C. Colonies were picked in a 96 deep well plate containing 400 µL of LB with kanamycin and chloramphenicol and incubated overnight at 37°C 220 rpm. A glycerol stock plate was made by using 25 µL of 87% glycerol and 75 µl of culture from each well. The plate was stored at -80°C. The remaining culture was used for Sanger sequencing (Microsynth Seqlab). After obtaining sequencing results, the desired strains were individually cultivated and stored at -80°C and the presence of their genomic mutation was asserted by using colony PCR as mentioned above.

### Construction of the double mutant Δ*punC*/*purA*^L75D^

JW1651-1 Δ*punC* from the Keio collection was used to construct the double mutant PurA^L75D^Δ*punC*. First, the kanamycin cassette of strain JW1651-1 Δ*punC* was cured by transforming pCP20 using electroporation^39^. Colonies were picked and cultivated for 20 h in LB medium at 43°C. Strains were then streaked on LB agar with no antibiotics and incubated overnight at 43°C. Multiple colonies were then picked and streaked on LB agar with either kanamycin, carbenicillin or no antibiotics, and incubated at 37°C overnight. Colonies with no resistance against carbenicillin and kanamycin and visible on LB agar were picked for colony PCR of the *punC* gene. PCR amplicons were sent for sequencing and strains harbouring the FRT scars and absence of the kanamycin resistance cassette were cultivated and stored at -80°C for further editing using the CRISPR-based genome editing method as described above.

### Construction of CRISPRi strains

Plasmids pgRNAK-purA#1 (protospacer: 5’-TTTACCTTCGTCACCCCATT-3’) and pgRNAK-mRFP (protospacer: 5’-AACTTTCAGTTTAGCGGTCT-3’) were constructed by exchanging the ampicillin resistance cassette of the original pgRNA plasmid^40^ with a kanamycin resistance cassette. Plasmids were then transformed into YYdCas9^37^ using electroporation.

### Time-kill assay

Precultures in 4 mL LB were inoculated from glycerol stocks for 8h at 37°C and 220 RPM and transferred to M9 medium for overnight incubation at 37°C and 220 RPM. M9 pre-cultures were used to re-inoculate shake flasks containing 25 mL of M9 medium with kanamycin and chloramphenicol. When OD600 0.25 was reached, 10 mL of medium were transferred to a new shake flask and carbenicillin was added at a final concentration 50 µg/mL. Cells were incubated at 37°C and 220 RPM. At each time point, 1 mL of the culture was sampled into a microcentrifuge tube and cells were centrifuged at 3000 RPM for 10 min. Supernatant was discarded and cells were resuspended in 1 mL of fresh M9 medium without carbenicillin. This washing step was then repeated. Cells were then serial diluted in fresh M9 medium by a factor of 10 to obtain dilutions of 1:100, 1:1000, 1:10000 and 1:100000. 100 µL from each dilution were then plated on M9 agar and incubated 48h at 37°C. Colonies were then counted to quantify colony forming units (CFUs) per mL. Time-kill assays with CRISPRi-*purA* and CRISPRi-*mRFP* strains was performed similarly.

For time-kill assays with clinical isolates, precultures of isolate #1 were first performed in LB for 7-8h at 37°C. Each preculture was then split into one M9 preculture with 100 µM adenine and one preculture without adenine, and grown overnight at 37°C. The next day, cultures in exponential phase were used to inoculate shake flasks with 25 ml of M9 medium with or without 100 µM adenine, with starting OD600 0.25. Sulbactam (TCI #S0868) was added at a final concentration of 12.5 µg/mL and carbenicillin was added at final concentration of 100 µg/mL. At indicated time points, 1 ml of culture were sampled from each flask and cells were washed with M9 medium supplemented with 5g/L glucose and 100 µM adenine, and plated on M9 agar supplemented with 100 µM adenine.

### Proteomics measurement

For proteomics, *E. coli* cells pellets were resuspended in 300 μL lysis buffer (0.5% sodium lauroyl sarcosinate (SLS), 100 mM ammonium bicarbonate) and heated for 15 min at 90°C. Proteins were reduced with 5 mM Tris(2-carboxyethyl) phosphine (Thermo Fischer Scientific) at 90°C for 15 min and alkylated using 10 mM iodoacetamid (Sigma Aldrich) at 20°C for 30 min in the dark. The totally extracted protein material was digested with 1 µg of trypsin (Serva) at 30°C overnight. After digestion, SLS was precipitated by adding a final concentration of 1.5% trifluoroacetic acid (TFA, Thermo Fischer Scientific). Peptides were desalted by using C18 solid phase extraction cartridges (Macherey-Nagel). Cartridges were prepared by adding acetonitrile (ACN), followed by equilibration with 0.1% TFA. Peptides were loaded on equilibrated cartridges, washed with 5% ACN and 0.1% TFA containing buffer and finally eluted with 50% ACN and 0.1% TFA, and finally dried. Dried peptides were reconstituted in 0.1% Trifluoroacetic acid and then analyzed using liquid-chromatography-mass spectrometry carried out on a Exploris 480 instrument connected to an Ultimate 3000 RSLC nano and a nanospray flex ion source (all Thermo Scientific). Peptide separation was performed on a reverse phase HPLC column (75 μm x 42 cm) packed in-house with C18 resin (2.4 μm; Dr. Maisch). The following separating gradient was used: 98% solvent A (0.15% formic acid) and 6% solvent B (99.85% acetonitrile, 0.15% formic acid) to 35% solvent B over 30 minutes at a flow rate of 300 nl/min. MS raw data was acquired on an Exploris 480 (Thermo Scientific) in data independent acquisition mode with a method adopted from ^41^. In short, Spray voltage were set to 2.0 kV and heated capillary temperature at 275 °C. For DIA experiments full MS resolutions were set to 120.000 at m/z 200 and full MS, AGC (Automatic Gain Control) target was 300% with an IT of 50 ms. Mass range was set to 350–1400. AGC target value for fragment spectra was set at 3000%. 45 windows of 14 Da were used with an overlap of 1 Da. Resolution was set to 15,000 and IT to 22 ms. Stepped HCD collision energy of 25, 27.5, 30 % was used. MS1 data was acquired in profile, MS2 DIA data in centroid mode.

Analysis of DIA data was performed using DIA-NN version 1.8 ^42^ with a *E. coli* uniprot protein database. Full tryptic digest was allowed with two missed cleavage sites, and oxidized methionines and carbamidomethylated cysteins. Match between runs and remove likely interferences were enabled. The neural network classifier was set to the single-pass mode, and protein inference was based on genes. Quantification strategy was set to any LC (high accuracy). Cross-run normalization was set to RT-dependent. Library generation was set to smart profiling. DIA-NN outputs were further evaluated using the SafeQuant^43,28^ script modified to process DIA-NN outputs.

### Whole genome sequencing of *E. coli* clinical isolates

DNA was extracted using the DNeasy UltraClean Microbial Kit (Qiagen), followed by library preparation (Illumina DNA Prep, (M) Tagmentation, Illumina) and barcoding (IDT for Illumina DNA/RNA UD Indexes). Sequencing was perform using a MidOutput Cartridge (NextSeq 500/550 Mid Output Kit v2.5 (300 Cycles)) on an Illumina NextSeq 500 machine.

### Growth experiment with clinical isolates

Cultivation of isolate #1 and #6 was performed as for the time-kill assay. OD600 was normalised for each precultures and strains were added to a 96-well plate containing fresh M9 medium with or without 100 µM adenine. The plate was then incubated for 24 h at 37°C in a BioTek Epoch plate-reader.

