## Supplementary Figures for "Purine nucleotide limitation undermines antibiotic action in clinical *Escherichia coli*"

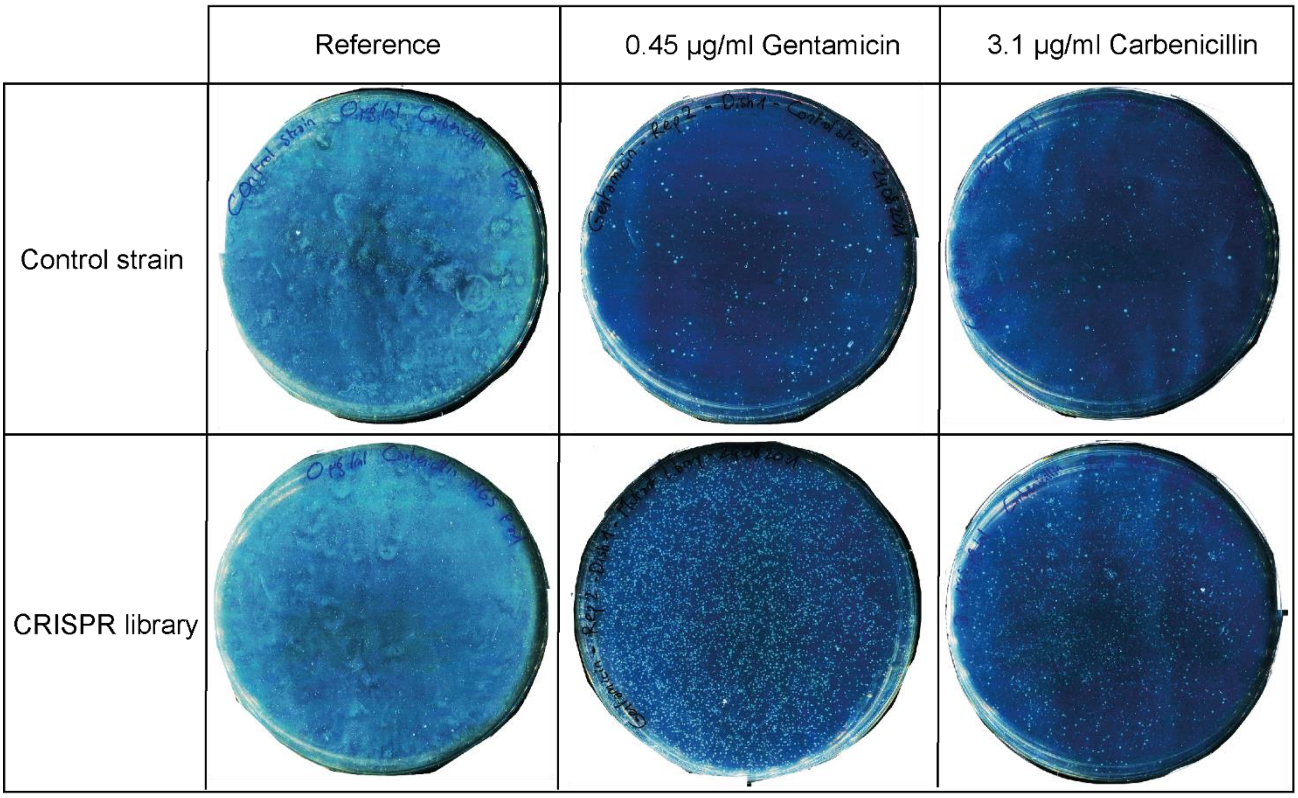


**Fig. S1.** Agar plates inoculated with the control strain (top row), or the CRISPR library (bottom row). Plates contained no antibiotics (reference, left column), gentamicin (middle column), or carbenicillin (right column).

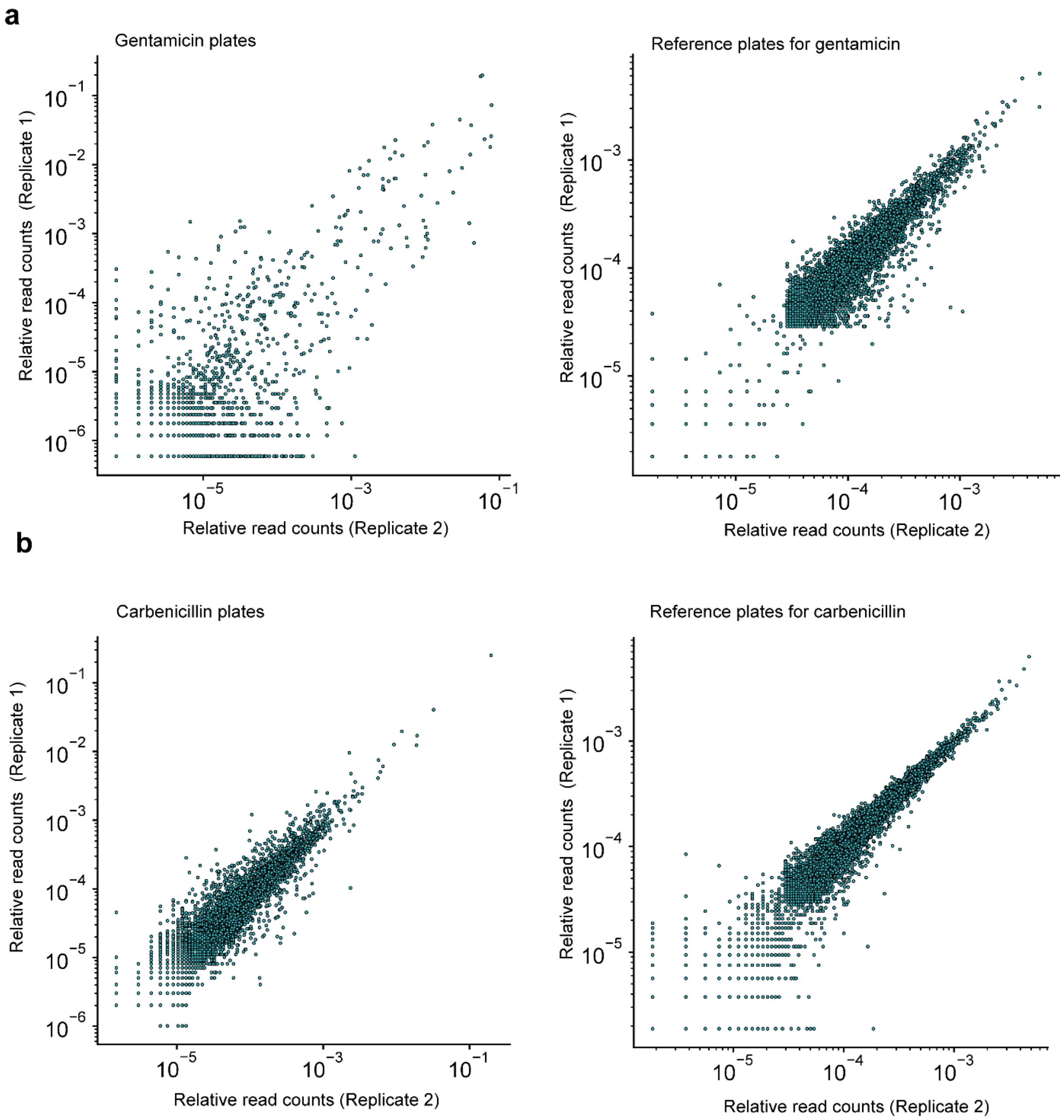


**Fig. S2**. a, Barcode abundances (relative read counts) on plates with gentamicin and the respective reference plates. b, Barcode abundances on plates with carbenicillin and the respective reference plates. Shown are the two replicates performed at different days.

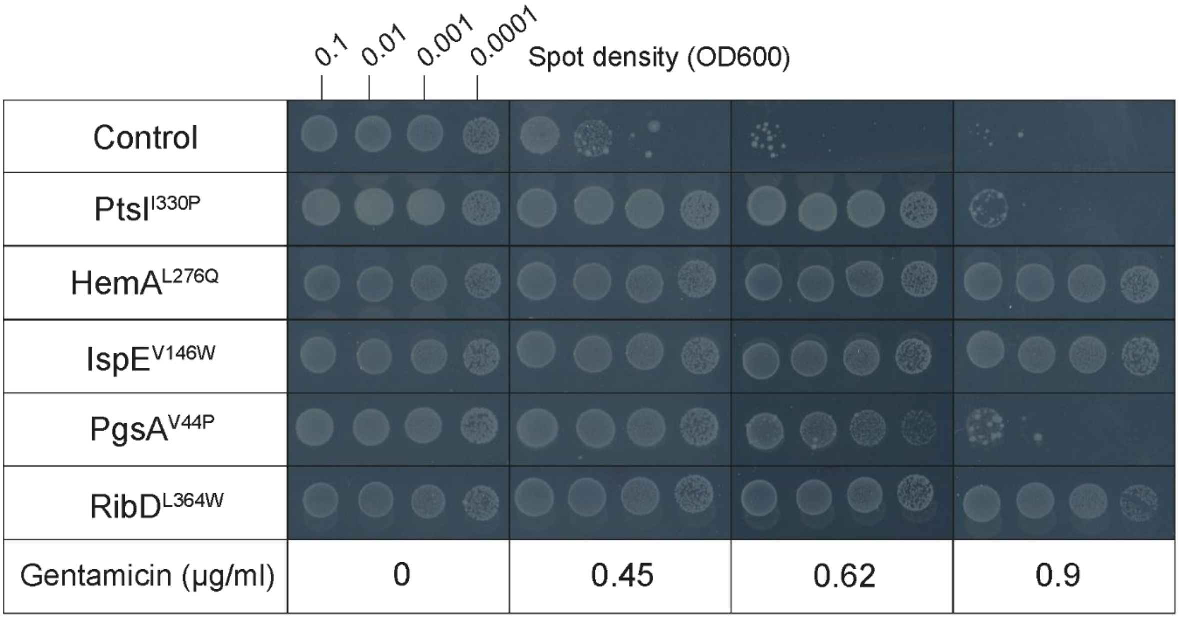


**Fig. S3.** Agar dilution assay with the control strain and gentamicin screen enriched mutants (PtsI$^{I330P}$, HemA$^{L276Q}$, IspE$^{V146W}$, PgsA$^{V44P}$, RibD$^{L364W}$). Each strain was spotted on agar plates with minimal glucose medium containing increasing concentrations of gentamicin (MIC = 0.45 µg/mL). Multiple inoculum densities were used to assess inoculum effects. Plates were incubated 48 h. Shown is one of n = 2 replicates.

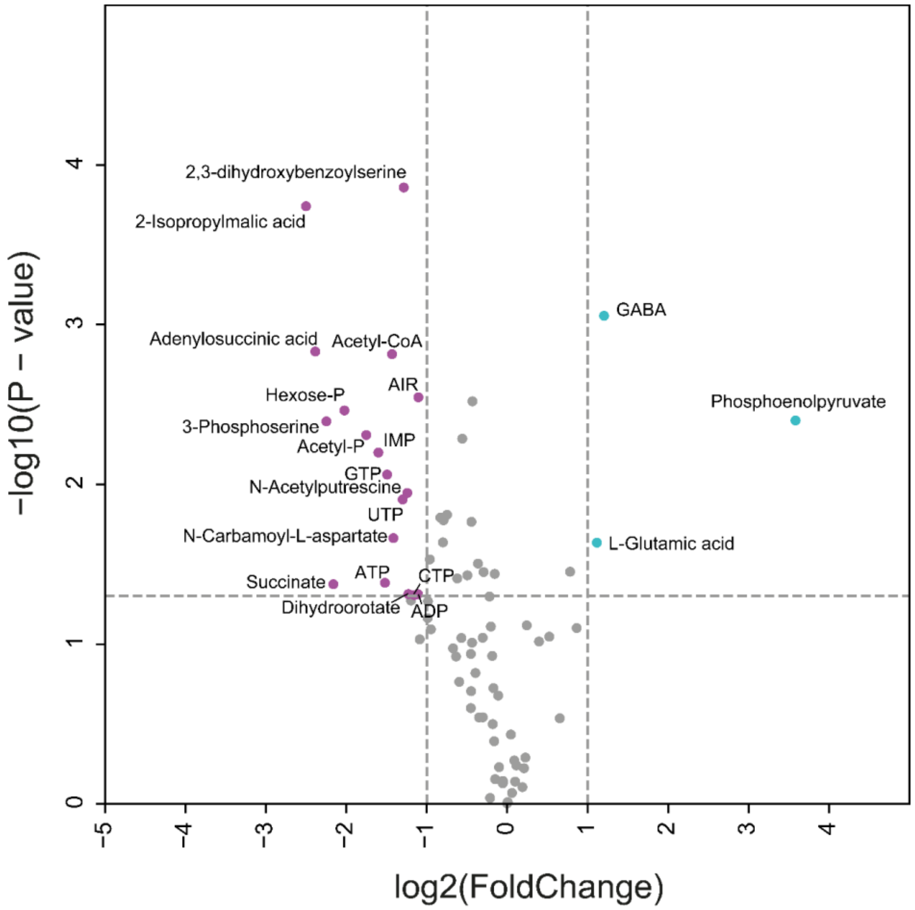


**Fig. S4.** Metabolome of the PtsI$^{I330P}$ mutant during drug-free exponential growth on glucose minimal medium. Shown are fold-changes of 82 metabolites relative to the un-edited control strain together with p-values. n = 3 samples from shake flask cultures.

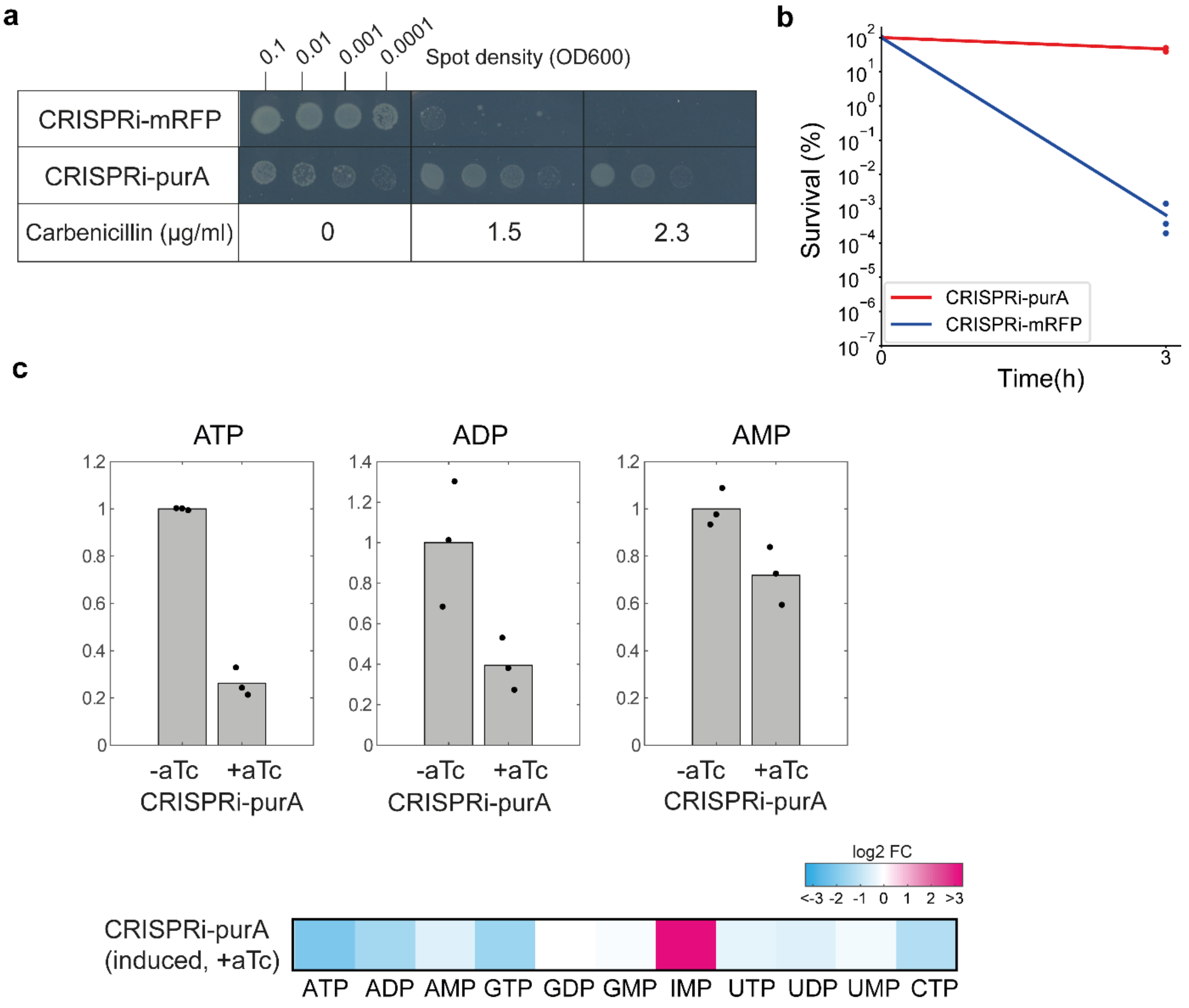


**Fig. S5.** a, Agar dilution assay with the CRISPRi-*purA* and CRISPRi-*mRFP* strains. Each strain was spotted on agar plates with minimal glucose medium and dCas9 inducer aTc (1 µM). Plates contained increasing concentrations of carbenicillin. Multiple inoculum densities were used to assess inoculum effects. Plates were incubated 48 h. Shown is one of n=2 replicates. b, Time-kill assay with the CRISPRi-*purA* and CRISPRi-*mRFP* strains. Each strain was incubated for 3 h with minimal glucose medium (0.2 µM aTc and 50 µg/ml carbenicllin). Survival shows colony forming units (CFUs) at the respective time point normalized to CFUs before drug exposure (t= 0 h). Dots show n = 3 replicates, and lines connect the mean. c, Nucleotide levels in the induced CRISPRi-*purA* strain during exponential growth with 0.2 µM aTc (+aTc). Shown are fold-changes of nucleotides relative to the un-induced CRISPRi-*purA* strain (-aTc).

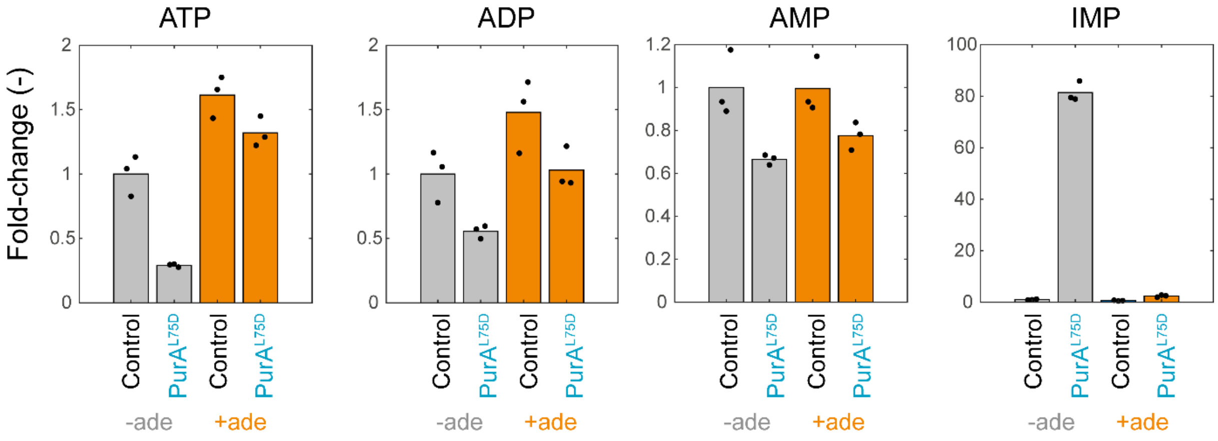


**Fig. S6.** Relative concentration of nucleotides (ATP, ADP, AMP and IMP) in the control strain and the PurA$^{L75D}$ strain. Strains were cultivated on minimal glucose medium with supplementation of 1 mM adenine (orange bars) and without (grey bars). Data are normalized to the control strain without adenine. Bars are the mean of n = 3 replicates (black dots). Data without adenine is the same as in Fig. 2 and shown as a reference.

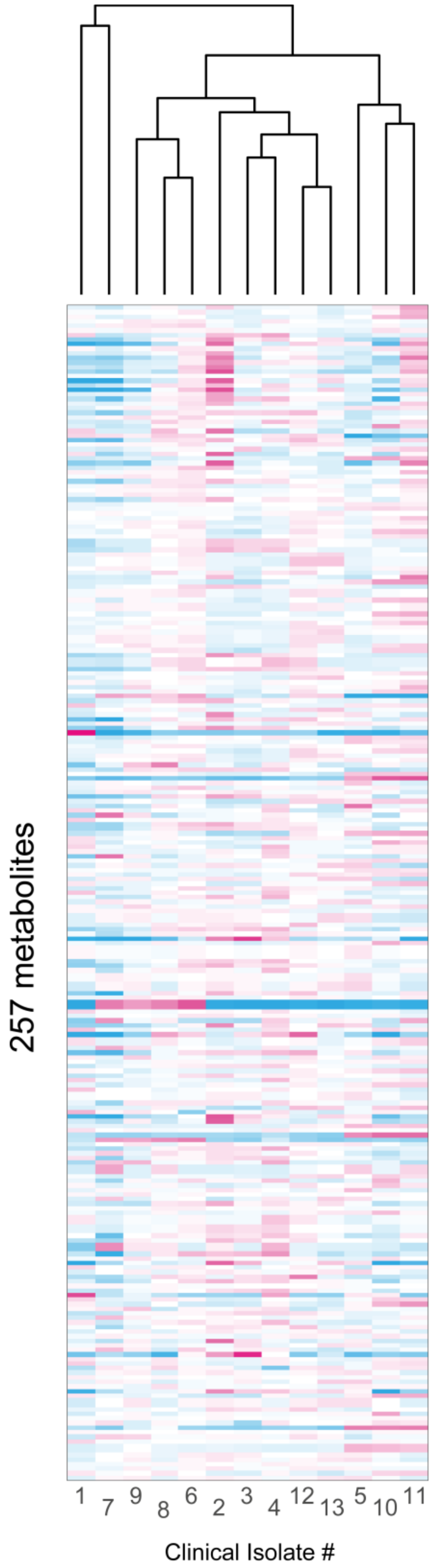


**Fig. S7.** Metabolome of 13 clinical *E. coli* isolates from urine of patients with suspected urinary tract infection. Shown are log2 fold-changes of 257 metabolites relative to the mean across all 13 strains. Only metabolites with a relative standard deviation lower than 50% in n = 3 replicates are shown. The dendrogram indicates the similarity of metabolome profiles.
